# Fast algorithm for determining orientations using angular correlation functions and Bayesian statistics

**DOI:** 10.1101/074732

**Authors:** Lanqing Huang, Haiguang Liu

## Abstract

Cryogenic electron microscopy (cryo-EM) method achieved revolutionary improvement in the past a few years, yet the orientation recovery for each particle remains computational challenging. Orientation determination can be solved using Bayesian approach, through which a model can be constructed to best match a whole set of experimental projections at their ‘correct orientations’. Without considering the centering of each particle projection, there are three degrees of freedoms (three Euler angles) that need to be fixed, usually resulting a computational complexity *O*(*n*^3^) where *n* is the number of discretization for each rotation angle. Here, we propose a method based on Maximum Likelihood approach with angular auto-correlation function of each projection, which is utilized to decouple the determination of three Euler angles to stepwise determination of two Euler angles and the subsequent third angle, the in-plane rotation. This approach reduces computational complexity from *O*(*n*^3^) to *O*(*n*^2^). Using simulation data, the accuracy and speed of the method is compared with the original maximum likelihood approach. We also investigated the impact of noise to the performance of this proposed method.

## 1 Introduction

Macro-molecular structure determination by single-particle analysis of electron cryo-microscopy (cryo-EM) images is a rapidly evolving field (Cheng et al., 2015; Cheng, 2015; Frank, 2016). Cryo-EM is a powerful structure determination method, using electron beams to “take photographs” of macromolecular complexes, such as viruses or protein complexes. The resolution of cryo-EM models has been improved to atomic resolutions, because of the three major breakthroughs: (1) sample preparation and handling that reduce the background noises; (2) the fast direct electron detection camera allows imaging in the movie mode that can correct the blur due to sample motion during exposure; (3) the algorithm development that enables model reconstruction and refinement at high accuracy within reasonable computing time. These developments have been reviewed in the recent special issue published in the journal of Microscopy (Vol 65, Issue 1).

The cryo-EM single particle reconstruction problem is challenging, because the molecules are embedded in noncrystalline ice at unknown orientations. To render these 2D experimental images into a high resolution three-dimensional model, one must use single particle 3D reconstruction approaches, which depends on accurate orientation determination. The computational breakthrough that allowed near-atomic reconstruction came from the development of statistical algorithms based on maximum-likelihood (ML) principles. The first application of maximum Likelihood method in single particle problem could be dated back to 1998 (Sigworth, 1998). Recently, a statistical framework for single particle cryo-EM data analysis was established where optimal parameters for model reconstructions could be inferred from the data without user intervention (Scheres, 2012a), and one successful implementation of this Bayesian framework is the RELION package by Scheres(Scheres, 2012b). RELION uses iterative approach to refine the projection orientations and the reconstructed model in turns. With this approach, several molecular complexes have been solved to near atomic resolutions from single particle cryo-EM data(Liao et al., 2013; Lu et al., 2014; Wong et al., 2014). Nevertheless, the computational cost for processing large amount of single particle projections is an obstacle preventing the methods to be used in a high throughput manner. For example, it is reported that approximately 100,000 core hours were required to process a dataset with 200,000 images and to obtain a refined 3D model using RELION package. Several strategies, such as domain reduction (Scheres et al., 2005) and grid interpolation (Tagare et al., 2008), have been implemented to reduce computational demands. The principle behinds such strategies is essentially the same: divide-and-conquer. Following this idea, we propose a *de novo* approach that significantly reduces computational time.

The original maximum likelihood approach computes the likelihood for each projection being in a set of reference projections by discretizing the 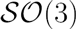 space spanned by three Euler angles ((*α*, *β*, *γ*)), therefore, it results a computational complexity of 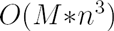, where *n* is the number of grid point for each Euler angle, and *M* is the number of experimental projection images. In contrast, the proposed approach recovers the orientation in two steps: (1) the electron beam direction relative to the molecule; and (2) the in-plane rotation of the resulted projection. This is achieved by transforming each projection image to an angular auto-correlation function (ACF). It can be shown that the ACF function does not depend on the in-plane rotation, because the auto-correlation function depends on angle difference ∆*γ*, rather than the rotation angle *γ*. This allow one to determine the first two Euler angles (*α*, *β*) using the auto-correlation function *C*(*r*, ∆*γ*), and then subsequently find the best matched in-plane rotation *γ* using the original projection (See Method section for details).

To evaluate the performance of the proposed stepwise orientation determination (SOD). We generated simulation data and tested the SOD method at various noise levels. The results show that the SOD method can recover the orientations correctly for cases with low noise levels. The speed-up is significant compared to the conventional maximum likelihood approach. When the noise level is high, the accuracy of the method is reduced, probably because the noise is amplified during the angular auto-correlation function calculation process. An advanced scoring function needs to be derived to enhance the noise tolerance. We also demonstrated that the same method can be applied to the orientation determination for single particle X-ray scattering data.

## 2 Methods

### 2.1 Implementations of Maximum Likelihood Approach

The detailed implementation of maximum likelihood approach is summarized by Scheres (Scheres, 2012b). Here, we provide a brief description. In the Bayesian approach, the reconstruction problem has been formulated as finding the model with parameter set Θ, including noise variance *σ* and other information, that has the highest probability of being the correct set in the presence of both the observed data 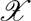 and the prior information 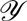. The variance 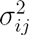 of the noise components is unknown and will be estimated from the data. Variation of 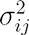 with resolution allows the description of non-white, or coloured noise. According to Bayesian law, the likelihood 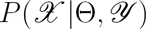 quantifies the probability distribution of variable 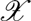 for given values of the hidden variables Θ. The expectation maximization algorithm is used to find the Euler angle (*α*, *β*, *γ*) and model class *k* (*k* = 1, if there is a single structure). For example, to find orientation variables (*α*, *β*, *γ*) for an experimental projection, the likelihood can be represented as (*k* is omitted as we are considering the simple case with *k* = 1):

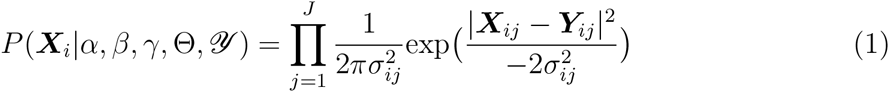

where 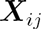 is value of *j*th pixel in *i*th experimental projection observed data 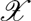.

In our implementations, Euler angle is defined by following the convention by the Heymann, Chagoyen and Belnap(Heymann et al., 2005): The first rotation is denoted by (*α*) or rot and is around the Z-axis; The second rotation is called (*β*) or tilt and is around the new Y-axis; The third rotation is denoted by (*γ*) and is around the new Z axis.

### 2.2 Angular auto-correlation Function, ACF

Kam proposed an ‘angular correlation’ function for the analysis of X-ray scattering patterns resulted from multiple molecules(Kam, 1977). Here, this correlation function is utilized to decouple three Euler angles, so that the rotations that fixed the beam direction and the in-plane rotation angles can be determined stepwisely.

The angular auto-Correlation Function (ACF) is defined as:

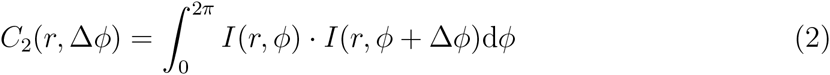

where *I*(*r*, *ϕ*) is the intensity of projection at a pixel position in polar coordinates (*r*, *ϕ*) on the detector.

The stepwise orientation determination (SOD) approach can be formulated as the following by computing two probabilities *P*_1_ and *P*_2_, with the latter only exams likely cases screened by *P*_1_:

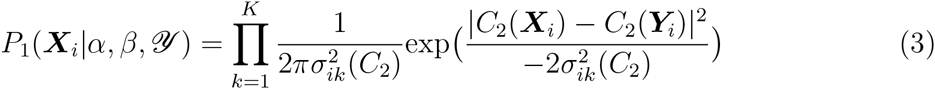

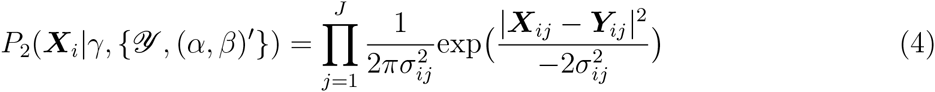

Where (*α*, *β*)′ is a subset that yields reasonable *P*_1_ values among all (*α*, *β*). Note that the 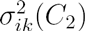 in equation (3) is the noise estimation of the ACF function. The computational complexity in this stepwise approach can be estimated as 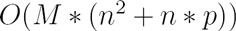, if we select the best matched *p* beam directions in the first step as plausible orientation candidates for the second step *P*_2_ calculations. When the signal-noise-ratio (SNR) is high, it is possible to identify a subset of *p* directions from *n*^2^ choices, such that the computational complexity can be written as 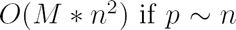. The extra overhead is the ACF calculation, which linearly depends on the number of experimental images *M*, and the number of reference images *n*^3^. This overhead is negligible compared to the product 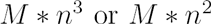.

### 2.3 Coordinate mapping methods

In order to compute the ACF from a projection image, the pixel values at position (x,y) needs to be mapped to polar coordinates (*r*, *ϕ*). The nearest neighbour mapping and the simple linear interpolation approach were compared.

The difference between any two images (projection image or the ACF image) is measured as the following:

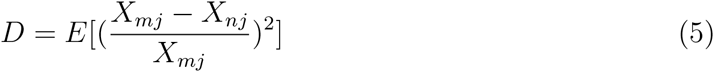

where *m, n* denote the image indices, and the subscript *j* = 1,…, *J* is the pixel indices of images *m* or *n*.

## 3 Experimental procedures and results

### 3.1 Rotational invariant property of ACF images

The ACF function has integrated out the azimuth angle *ϕ*, so the ACF image does not depend on the in-plane rotation angle *ϕ*. This is verified numerically, which is shown in figure 1, simulation projections at different *γ* angles with the same (*α*, *β*) correspond to the same ACF image.

**Figure 1:**
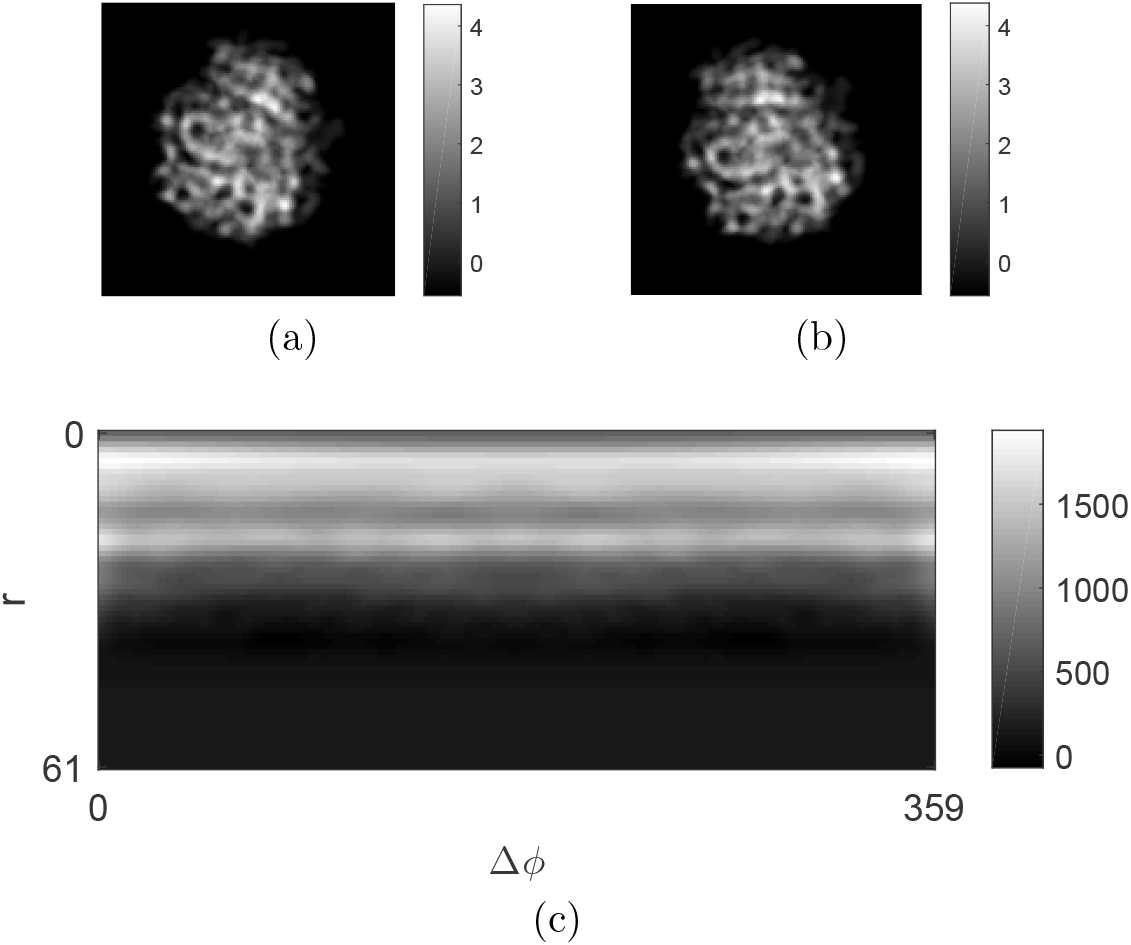
Correlation image does not depend on in-plane rotation. (a) The projection of particle EMD-6044 from Euler angle 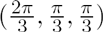. (b) The projection of particle EMD-6044 from another view 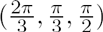. (c) ACF Image generated from above two particle projections.

The image difference quantity *D* is measured between an image and its own rotated counterparts. It is shown that in Figure 2a where the value *D* varies significantly while the image is rotated from 0 to 2*π*. On the other hand, ACF image is effectively invariant of the rotation. Compared to original images, difference between ACF under rotation and the ACF of the reference image is almost zero, as shown in Figure 2b (Note the scale difference of the Y-axes). In the same figure, it is shown that the linear interpolation approach is more accurate than the nearest neighbour mapping when converting the cartesian coordinate image to polar coordinate image.

**Figure 2:**
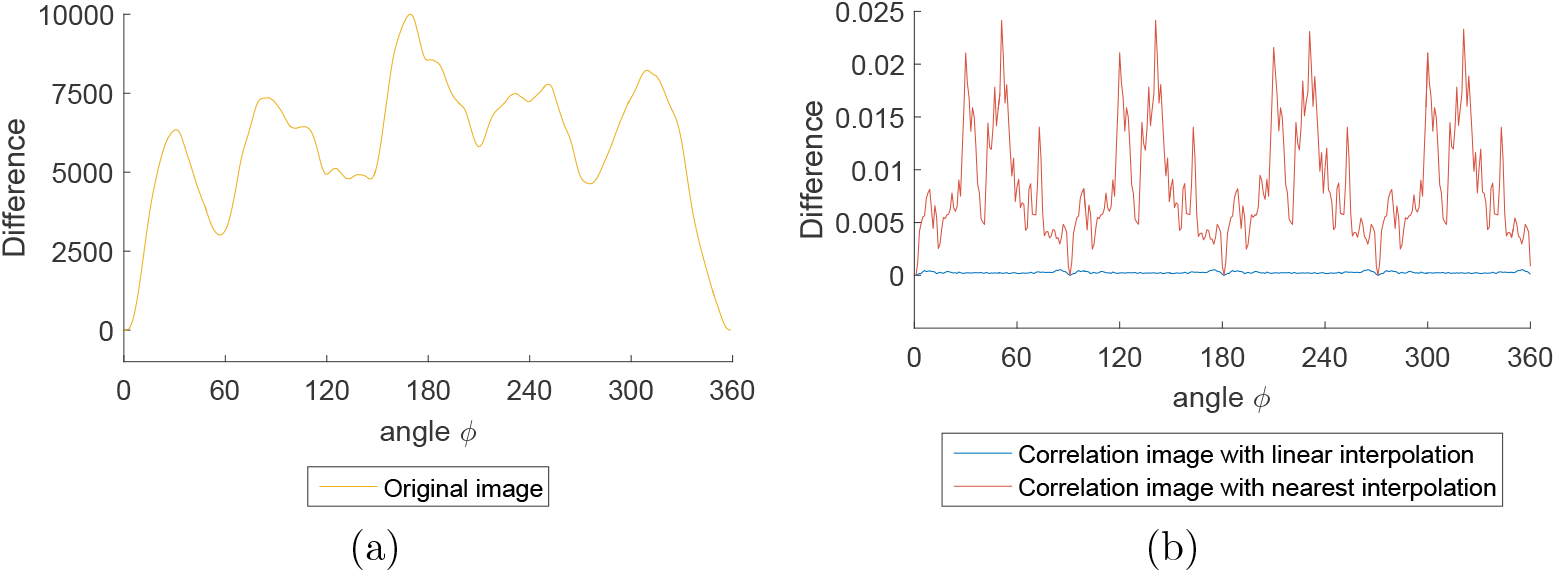
Rotational invariance of correlation image in real-space. (a) The difference of a set of projections in real-space from 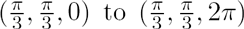 compared to a reference projection at orientation 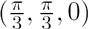. (b) The result of calculating difference for ACF image computed from the same projections in (a). Two interpolation approaches are compared, and the nearest neighbor approach is less accurate compared to the linear interpolation approach.

This lays the foundation for the new method in solving orientation problem. Two Euler angles will be determined using ACF function (see equation (3)). Then third Euler angle is found by Maximum-Likelihood method in a subsequent step using the original projection image (see equation (4)).

### 3.2 The ACF image of scattering patterns in Fourier space

The real space projection has another two parameters to fix the center of the image. In this work, we did not attempt to develop new method to determine the shifting parameters. We further explored the ACF rotational invariant property in Fourier space, where the shifting in X-Y plane is absorbed in the phase terms of the fourier transforms. The modulus of the fourier transform corresponds to scattering pattern intensity, which does not depend on the X-Y shift in real space.

We found that the ACF method applies to intensity images in fourier-space (see figure 3). Clearly, rotational invariance of ACF image perform well in fourier-space. Linear interpolation in fourier-space is slightly better than nearest neighbour mapping.

**Figure 3:**
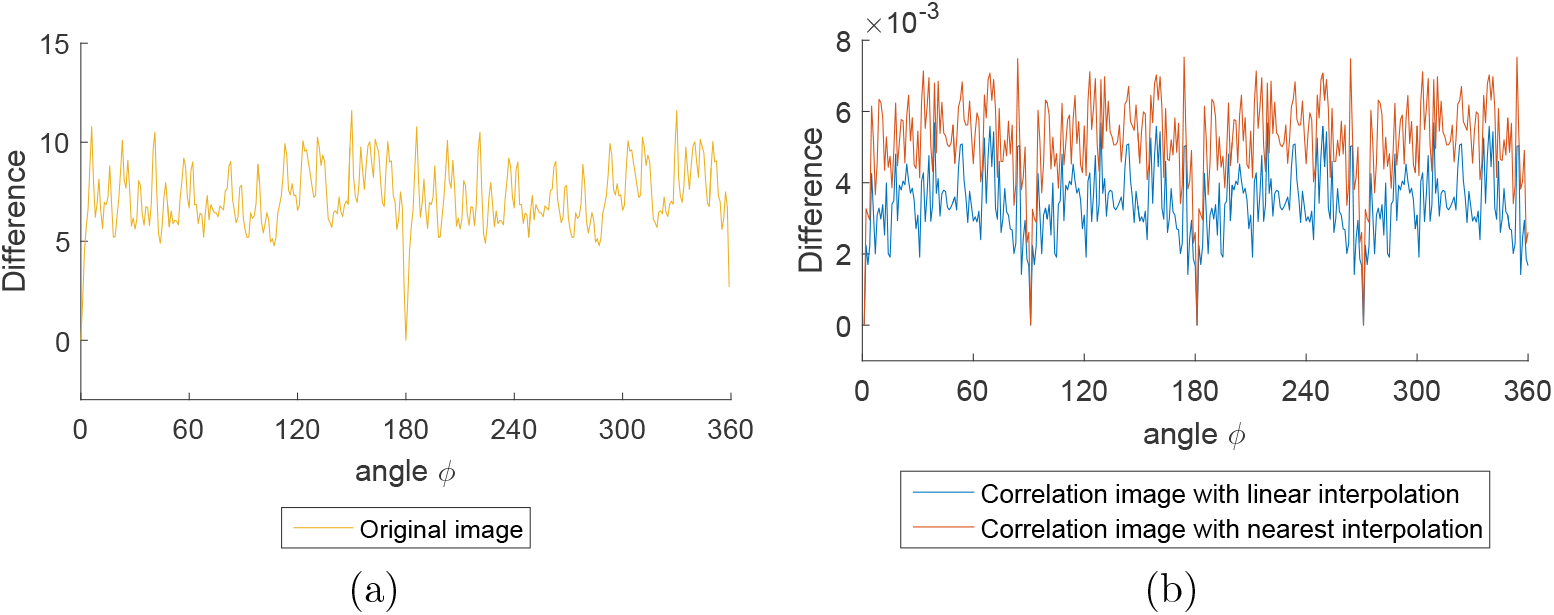
Rotational invariance of correlation image in fourier-space. (a) The difference of a set of projections in fourier-space from 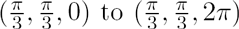 compared to a reference projection at orientation 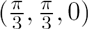. (b) The difference of a set of correlation images from 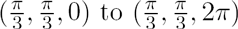 compared to a reference projection at orientation 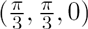. Note the value, indicating that the differences in correlation functions can be neglected.

It is noteworthy to mention that the periodicity observed from the different curves in both fourier and real space that is caused by interpolation errors in projection. An image which rotate every 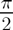 degree around *γ* can be exactly mapped from the original image without any interpolation. Therefore, we observe minimum difference values at these angles.

Without specifications, linear interpolation method is used in the analysis presented in this work.

### 3.3 Reducing computational error in fourier-space: oversampling

When transforming the real space projection image to Fourier intensity, the points in fourier space can be sampled at higher frequencies. This is equivalent to X-ray/electron single molecule scattering experiment with a setup that allows collecting data at a frequency higher than the value given by Shannon-nyquist sampling theorem. It is observed that the oversampling rate can affect the accuracy of interpolation.

Figure 4 gives the influence of difference value of image with different oversampling rates. From figure 4a and figure 4b, it is clear that larger oversampling rate is useful to reduce computational errors at the cost of longer computing time (Figure 4c). The improvement of accuracy can be attributed to finer sampling of intensity that reduces linear interpolation error. To balance the computing efficiency and accuracy, an oversampling rate of 5 was used in the transformation of projection images to Fourier intensity images.

**Figure 4:**
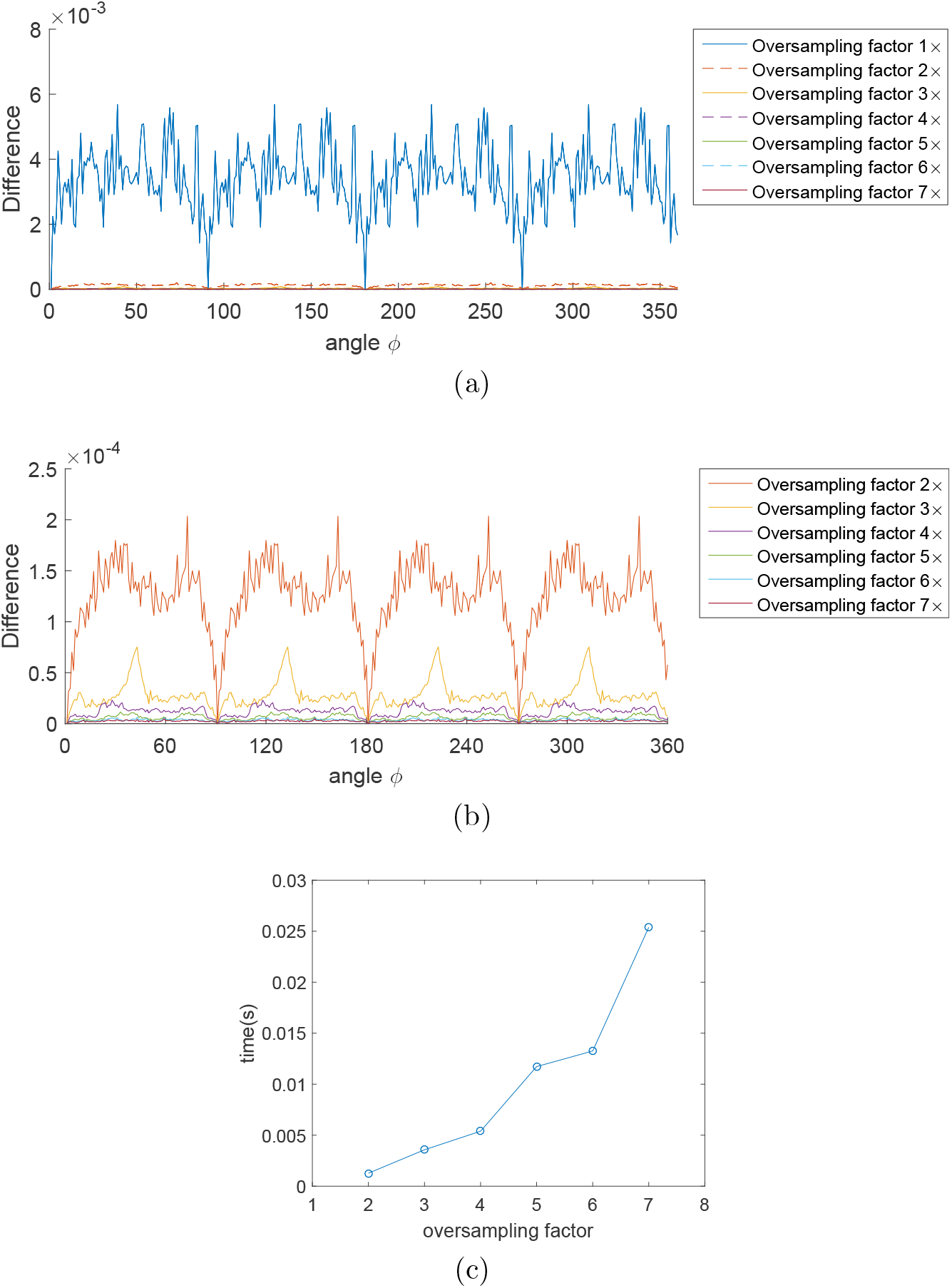
Reducing computational error by oversampling. (a) Each line in the figure is the result of calculating difference for a set of correlation images in fourier-sapce 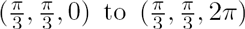 with a reference correlation image of 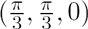 from 3D model (EMDB-6044). And oversampling factor changes from 1 to 7. (b) The detail information of difference for correlation image with oversampling factor 2~7. (c) Average time cost to oversample a image with different factors.

### 3.4 Improving robustness: adaptive method

If the single best matched (*α*, *β*) angles were used for the subsequent search of *γ*, the algorithm is very limited to high signal-noise-ratio situations. This does not fully utilize the power of maximum likelihood algorithm. Yet, the matching scores for all (*α*, *β*) combinations provide valuable information to screen out a large fraction of bad orientation candidates. The question is how to choose the *p* number of ‘good’ candidates.

The following method is used to select the plausible candidates: first the scores are ranked, and then the numerical difference between adjacent elements is calculated to estimate the derivatives, as shown in figure 5c. The derivatives are used to set the cutoff and the top *p* beam directions are further pursued for the fixation of in-plane rotation. In this study, the cutoff is found by locating the largest derivatives values.

**Figure 5:**
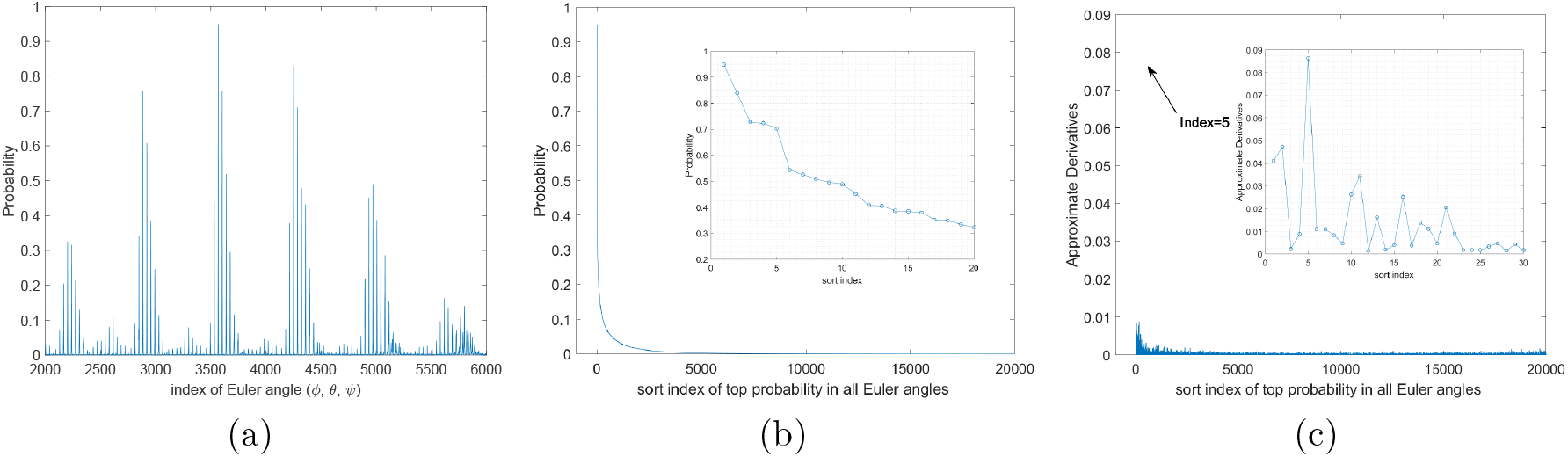
Process to output nearby angle. (a) is the original probability calculated by formula (1) or (3). (b) is sorted result of curve in (a). After sorting, approximate derivatives (figure(c) is calculated by derivatives) to find the maximum point which indicates the boundary in latent correct angles.

### 3.5 Accuracy performance under Poisson noise

The recording process, whether by means of a photographic plate or by a direct image pick-up device, is burdened with another source of noise, which is due to the quantum nature of the electron: the shot noise. this noise portion is caused by the statistical variations in the number of electrons that impinge on the recording target, which follows Poisson statistics.(Frank, 2006).

We used simulation data to quantify the influence of poisson noise to the performance of correlation function method. In the simulation process, Poisson noise were added to image to simulate experimental projection. Particle EMD-6044 (Dashti et al., 2014) was used to generate simulated experiments. In all cases the particle images provided were 125 × 125, and had a resolution of 3.0Å(0.30*nm* per pixel). In real-space, Poisson noises was added into projections of 3D model. For simulation projections of fourier-space, noise added to image in real-space and then transformed images to fourier-space. For Poisson noise distribution, the variance at pixel *j* for image *i* is set to be 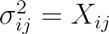.

#### The stepwise maximum likelihood method can recover orientations on reference grids

In this test, experimental projections were simulated form 3D model (EMDB-6044) with Euler angle on pre-generated grids. Experimental testing projections were divided into two classes. In our algorithm, an ACF image with specific rot and tile angles (*α*, *β*) can be used to represent for a series of particle images with same (*α*, *β*) angles but different in-plane rotation angles *γ*. For this reason, the particle projection with in-plane rotation *γ* = 0° was used generated the ACF image references. Hence, the distribution of angles in Class 1 was generated by random *α*(rot) and *β*(tilt) value with *γ* = 0. And angles in Class 2 randomly distributing in whole angle space (*α*, *β*, *γ*) with *γ* ≠ 0.

Figure 6 illustrates the accuracy performance of the proposed correlation based determination approaches with Poisson noise. SOD approach showed excellent performance and achieved the same accuracy compared to the Maximum Likelihood approach. Besides, comparing to class 2, class 1 does not present a better accuracy. It indicates that ACF image has a well defined rotational invariant property. In addition, those samples failed in find correct orientation fell in the cases with tilt angle *β* = 0 or *π*. Overall, for the Poisson noise taken by experimental devices, SOD approach gives a good performance.

**Figure 6:**
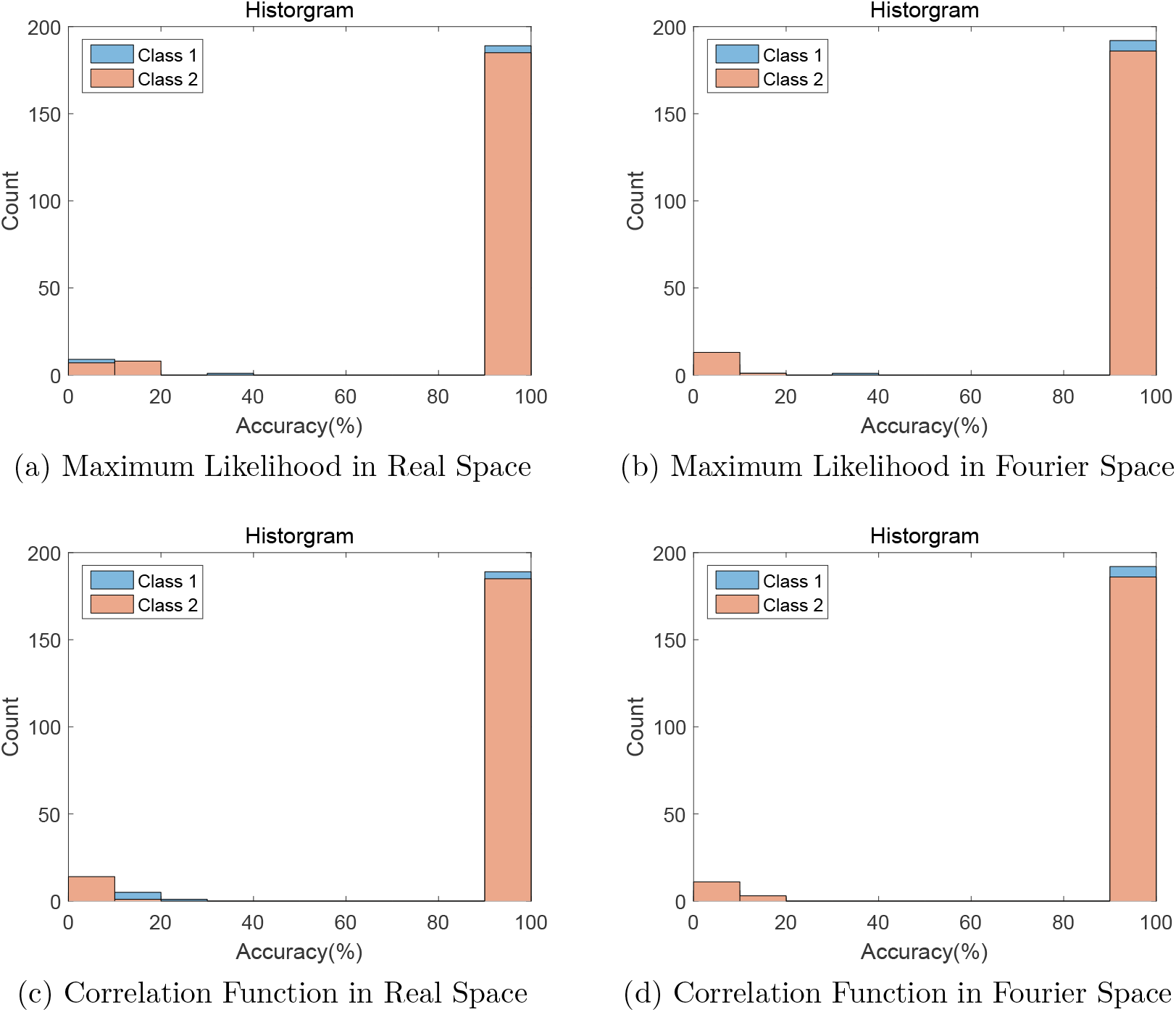
Accuracy performance histograms of different approaches with Poisson noise. (a)(b)(c)(d) are respectively corresponding to different approaches in different space. Reference projections came from 3D model of EMDB-6044. Grid size of references was equal to 10°. Both two classes included 200 random input samples and each sample repeated 100 times with different Poisson noise.

### 3.6 Accuracy performance for orientations that deviate from grid points

In real situations, experimental image does not have to be on the reference sampling grid. Hence we need to know how well the SOD approach performs in the cases where the projection orientations deviated from reference grids. Here we tried to determine orientations for projections that are off the reference orientation grids at different levels, with Poisson noise situation in section 3.5.

Figure 7 presents correlation approach works well in the cases where the simulated projections deviate from pre-defined reference orientation grids. It is interesting to observe that the SOD approach performs better than the conventional Maximum Likelihood method.

**Figure 7:**
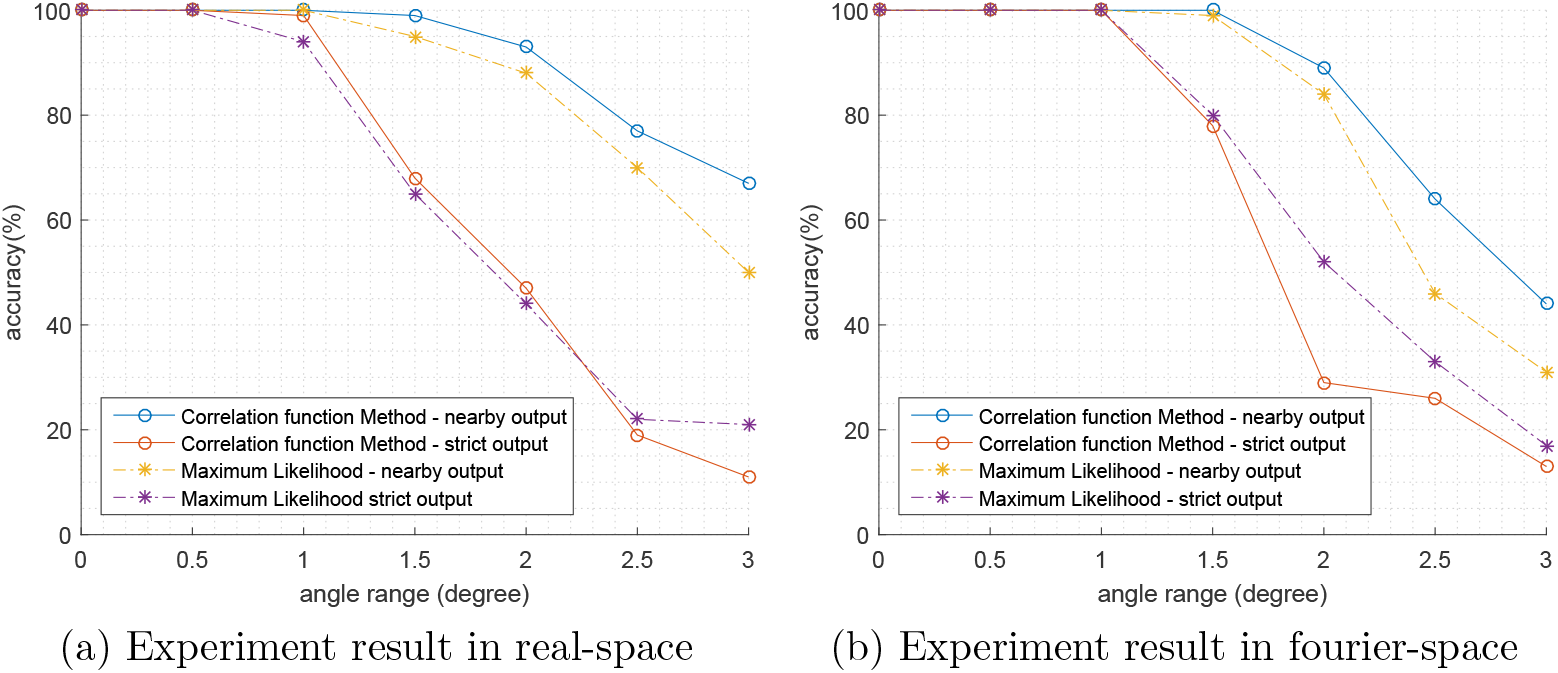
Perturbation accuracy performance of different approaches. X axis stands for angle range of samples deviated from reference grids. Each angle range has 100 samples which were random projections of different Euler angles except for *β* = 0, π with Poisson noise from 3D model EMDB-6044. And all tests were based on the reference sampling grids equal to 3°.

### 3.7 Accuracy performance for images with Gaussian noise

In actual experiments, Gaussian noise plays a major role in particle projections mainly due to the background signals. To simulate Gaussian noise, original projections from 3D model (EMDB-6044) were created and pixel value of these projections image was scaled to the mean values of 0 and variance *σ*(*s*) = 1. Then, white noises with different variance *σ*^2^ followed Gaussian distribution *N*(0, *σ*) were added into original projections to generated actual experimental projections. Hence, Signal-to-Noise Ratio (SNR) was easy to calculated by

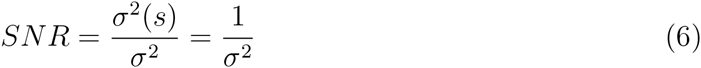

We assumed that ACF images (see Figure 1) still follow the Gaussian distribution. Therefore in equation 3, according to equation 2, an estimated value of *σ*^2^(*C*_2_) was approximated to be < *s*^2^ >_*ϕ*_ ·(*σ*^2^).

To test the performance of ACF algorithm under different SNR situations, 400 simulated experimental projection with random Euler angles (*α*, *β*, *γ*) with a specific noise level *σ* were generated, and both SOD and Maximum likelihood approaches were applied to find their correct orientations. Figure 8 shows results at different SNR levels. The accuracy of SOD algorithm under Gaussian noises decreased after SNR gets below 0.4.

**Figure 8:**
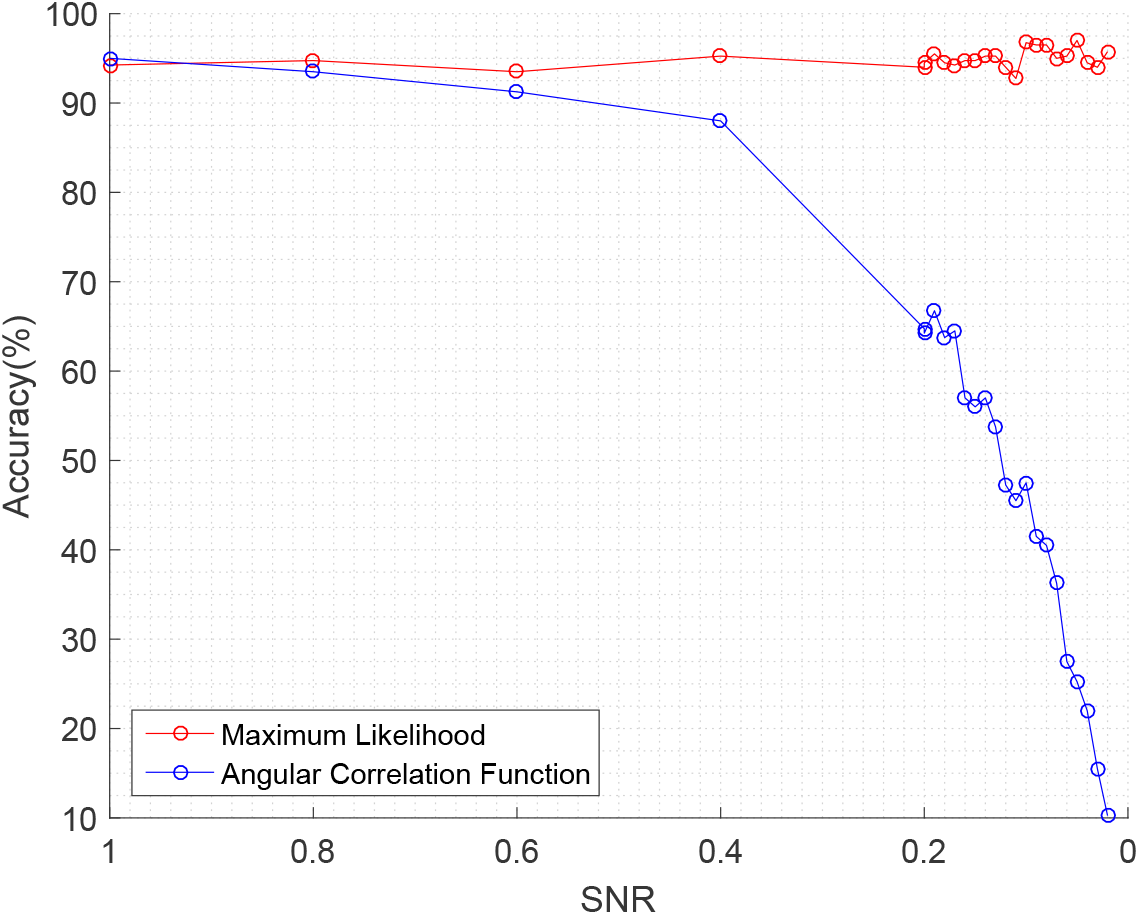
Accuracy performance of Gaussian noise. X axis stands for the signal noise ratio (SNR) of the simulated experimental projections. Y axis is the accuracy performance of orientating 400 simulated experimental projections with specific white noise created from particle EMD-6044. The step of reference grids was equal to 10°, and about 24,624 reference projections were created as prior information.

At present, particle projection images obtained in cryo-EM experiment often has poor Signal-to-Noise Ratio (SNR). Given the recent advancement of electron detector (DDD) cameras(Cheng et al., 2015), better quality data with good SNR is expected. On the other hand, more sophisticated scoring functions for SOD are desired to distinguish ‘good’ orientation candidates from the rest.

### 3.8 The stepwise orientation recovery significantly reduces computing time

The major advantage of the SOD method is that the de-coupling three Euler angles enables a boost of computational speed. Table 1 shows the time cost to find correct Euler angle (*α*, *β*, *γ*) of one experimental image in the prior information 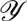. With the increasing number of reference samples, the effect of reducing time cost by correlation function becomes more pronounced, as the computational complexity *O*(*M · n*^2^) predicted.

**Table 1:**
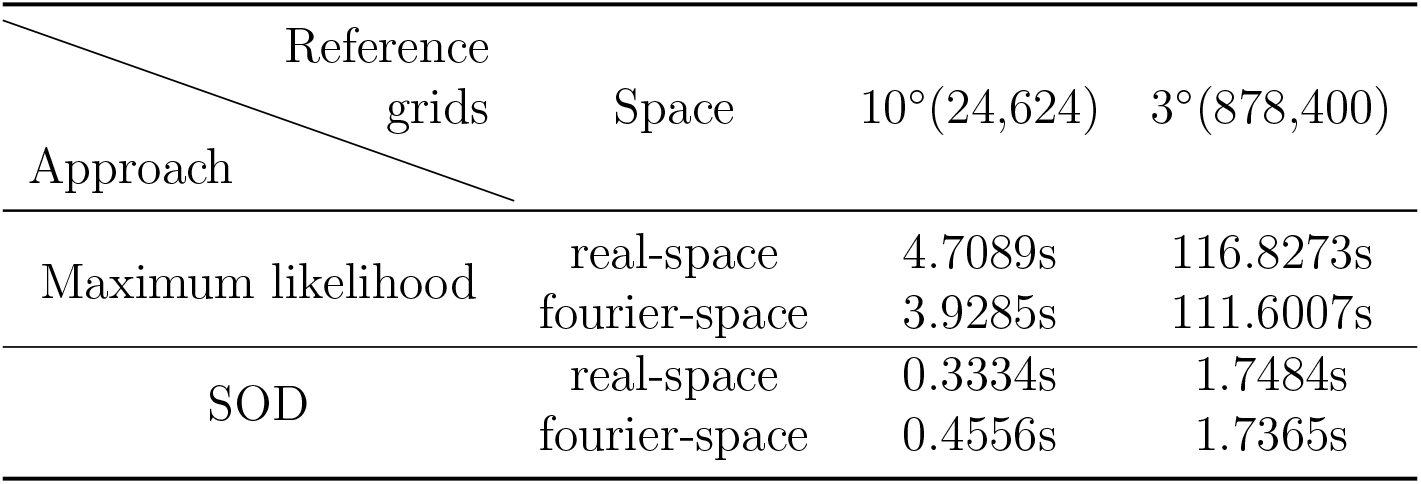
Average computational time for two approaches. Programs implemented by MATLAB run with the same parallel environment with 10 physics cores of intel E5-2600 v2 CPU. The numbers in parenthesis are the sizes of references dataset in the prior information 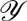.

Correlation function calculation of reference samples and experimental data is considered as overhead and has computational complexity of *O*(*M*) where *M* is the number of experimental/simulation images to be processed. In each maximization iteration, these correlation functions are calculated once and saved in memory for pairwise comparison. Therefore, this portion of computational cost is not significant compared to the overall required time.

## 4 Discussion and Conclusion

The computational complexity of the maximum likelihood algorithms sets the required computing time in the orientation recovery analysis. Although ‘divide-and-conquer’ approaches can be used to implement multi-level or multi-domain programs to reduce computing time, the overall computational complexity is not fundamentally reduced. In the proposed stepwise orientation determination approach, the degree of rotational freedom is reduced by decoupling the in-plane rotation from the other two rotations. Using simulation data, we have demonstrated the performance of the SOD algorithm and the speedup in computing. There are still two issues to be resolved to make the algorithm useful in real data analysis, namely, (1) the X-Y shifting problem, and (2) the analysis of data with low SNR’s.

The X-Y shifting problem is also known as the centering of projection images. This is critical for the proposed SOD algorithm, because the ACF image will not be correct without an accurately determined center for image conversion from cartesian coordinate to polar coordinates. In maximum likelihood approach, the shifting parameters are determined together with the rotation parameters in a combinatory manner. In the SOD approach, we propose to find the shifting parameters in an outer layer of the orientations. Specifically, for a given center, the likelihood of orientations are evaluated and saved for comparisons with the values calculated for different image centers. The most likely orientations and shifting parameters are determined using either a brutal force scanning of (*x*, *y*) around an initial estimated image center, or using some sort of optimization approach to refine the image center. Given the fact that 2D image classification and alignment can provide reasonable image centers, the local scanning or refinement of the X-Y shift should be doable.

The performance of SOD approach is not satisfactory for the cases that the image data has low SNR’s. We attribute this reduced accuracy to the noise amplifications during the ACF calculation. Furthermore, the error estimation for the ACF function is not straightforward, as the Gaussian noise coupling length is not available. In order to improve the accuracy of the likelihood evaluations, an improved error model has to be derived. We will pursue for such models to make the SOD approach applicable for the analysis of low SNR data.

As a summary, preliminary testing results show that the stepwise orientation determination approach works well in reducing the computational costs while maintaining the orientation recovery accuracy for the cases with SNR higher than 0.4. Robust algorithms and scoring functions need to be developed to make the method practical to handle actual experimental data with low signal-noise-ratio.

